# Hydrodynamic and Polyelectrolyte Properties of Actin Filaments: Theory and Experiments

**DOI:** 10.1101/2022.01.11.475973

**Authors:** Ernesto Alva, Annitta George, Lorenzo Brancaleon, Marcelo Marucho

## Abstract

Actin filament’s polyelectrolyte and hydrodynamic properties, their interactions with the biological environment, and external force fields play an essential role in their biological activities in eukaryotic cellular processes. In this article, we introduce a unique approach that combines dynamics and electrophoresis light scattering experiments, an extended semiflexible worm-like chain model, and an asymmetric polymer length distribution theory to characterize the polyelectrolyte and hydrodynamic properties of actin filaments in aqueous electrolyte solutions. A fitting approach was used to optimize the theories and filament models for hydrodynamic conditions. We used the same sample and experimental conditions and considered several g-actin and polymerization buffers to elucidate the impact of their chemical composition, reducing agents, pH values, and ionic strengths on the filament translational diffusion coefficient, electrophoretic mobility, structure factor, asymmetric length distribution, effective filament diameter, electric charge, zeta potential, and semiflexibility. Compared to those values obtained from molecular structure models, our results revealed a lower value of the effective G-actin charge and a more significant value of the effective filament diameter due to the formation of the double layer of the electrolyte surrounding the filaments. Contrary to the data usually reported from electron micrographs, the lower values of our results for the persistence length and average contour filament length agree with the significant difference in the association rates at the filament ends that shift to sub-micro lengths, the maximum of the length distribution.

## I. INTRODUCTION

Actin filaments (F-actins) are highly-charged double-stranded rod-like polyelectrolytes formed by the polymerization of G-actin proteins. Cytoskeleton filaments are essential for various biological activities in eukaryotic cellular processes. These filaments are usually organized into higher-order structures, forming bundles and networks which provide mechanical support, determine cell shape, and allow movement of the cell surface, thereby enabling cells to migrate, engulf particles, and divide. One major challenge in biophysics is to elucidate the role of the polyelectrolyte properties of the filaments, their interactions with the biological environment, and external force fields on their higher-order structure formation and stability. Indeed, it is imperative and crucial for understanding emergent or macroscopic properties of these systems. During the last few decades, a substantial amount of research has been done on the diffusion coefficient, shear modulus, second virial coefficient, and electrophoretic mobility of actin filaments[1–10]. Nevertheless, the underlying biophysical principles and molecular mechanisms that support the polyelectrolyte nature of F-actins and their properties still remain elusive. Sometimes, this uncertainty is due to the lack of unicity, consistency, and accuracy in the methodologies, techniques, and sample preparation protocols used in scattering experiments to produce meaningful, reproducible results. At the same time, the optimization of actin filament and electrolyte models and sophisticated molecular-level kinetic theories that characterize these macroscopic properties in hydrodynamic conditions became burdensome due to the use of parameters obtained in non-hydrodynamic (usually microscopy) conditions.

Nowadays, modern Dynamic Light Scattering (DLS) and Electrophoresis Light Scattering (ELS) instruments are robust and accurate tools to characterize hydrodynamics properties of polydisperse charged biomolecules even at low concentrations and using small sample volume. These non-invasive, susceptible, and resolution instruments use advanced technology and multi-functional software to measure the translational diffusion coefficient, second virial coefficient, and electrophoresis mobility with high accuracy and reproducibility. DLS and ELS experiments also allow for accurate measurement of model parameters if the number of these parameters is small and the approach is adequate to characterize the hydrodynamic and polyelectrolyte properties of the biomolecules in solutions[11, 12].

In this article, we introduce a unique approach that combines light scattering experiments and optimized theoretical approaches to characterize actin filaments’ polyelectrolyte and hydrodynamic properties. We used Malvern ULTRA Zetasizer instrument to measure actin filament’s translational diffusion coefficient and electrophoretic mobility at low protein concentration. We developed a novel sample preparation protocol based on bio-statistical tools[13] to minimize errors and assure reproducibility in our results. This protocol was used for all the experiments. We considered three different buffers, g-actin and polymerization, used in previous works[8–10] to elucidate the impact of their chemical composition, reducing agents, pH values, and ionic strengths on the filament properties. We also performed protein dialysis[14] and spectrophotometric[15] techniques to measure the protein concentration in our samples.

Additionally, we used a novel multi-scale approach to calculate the translational diffusion coefficient and electrophoretic mobility of polydisperse actin filaments in aqueous salt solutions. The monodisperse translational diffusion coefficient calculations are based on the Stokes-Einstein formulation[16] and a modified wormlike chain (WLC) model for the hydrodynamic radius[17]. The monodisperse electrophoretic mobility calculations are carried out using a linear polymer representation of the WLC, which accounts for the balance between forces acting on each chain’s monomer. This model and the Routine-Prager tensor for hydrodynamic interactions calculation are used to take the orientational average over all possible polymer conformations in the low electric field approximation[18]. An asymmetric, exponential length distribution is used to characterize the actin filament polydispersity and the different increasing rate lengths of barbed and pointed ends[19]. We used the length distribution to take the filament length average over the monodisperse translational diffusion coefficient and electrophoretic mobility expressions. The resulting expressions for the polydisperse translational diffusion coefficient and electrophoretic mobility depend on the persistence length, the effective filament diameter, the monomer charge, and the increasing rate length of barbed and pointed ends of the filaments. We used Mathematica software, a fitting approach, and multi-core computers to find optimal values for these parameters that better reproduce the translational diffusion coefficient and electrophoretic mobility values obtained experimentally.

### THEORETICAL APPROACH

#### Diffusion Theory

The Stokes-Einstein formulation provides the following expression to calculate the monodisperse translational diffusion coefficient of colloidal particles of any shape

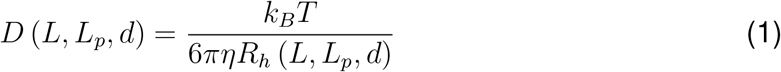

where *k_B_*, *T*, and *η* represent the Boltzmann constant, temperature, and viscosity of the dispersant, respectively. The hydrodynamic radius, *R_h_*, is also a factor in eq. (1) which depends on the contour length, *L*, persistence length, *L_p_*, and diameter of the filament, *d*.

The hydrodynamic radius is calculated using Mansfield’s approach for transport properties of semiflexible polymers[17]. The approach is based on the orientational preaveraging approximation. The charge distribution over the surface of an arbitrary shaped charged conductor is proportional to the Stokes-flow force distribution over the surface of a rigid body of the same size and shape as the conductor. Additionally, a cylindrical model of the WLC is used to account for not only the persistence, *L_p_*, and contour length, *L*, but also the diameter of the chain, *d* = 2*a*. (see Figure 1). As a result, the expression for the hydrodynamic radius of a semiflexible polymer is given by

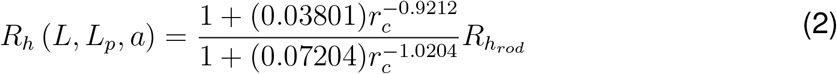

where

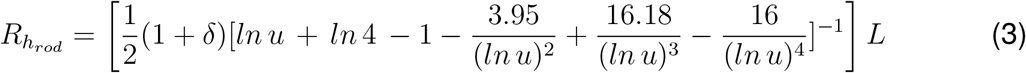

is the corresponding expression for a rigid polymer, and

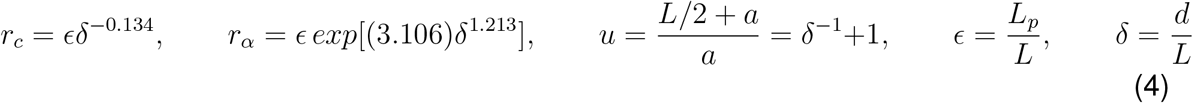

**Figure 1:**
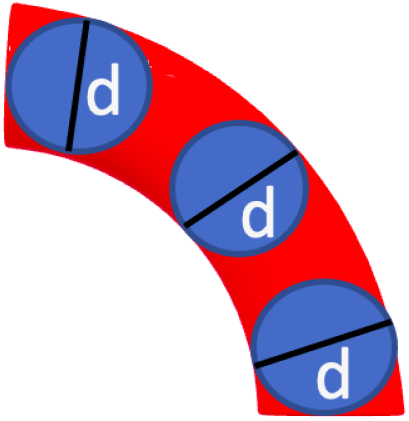
Cylindrical model of the wormlike chain enclosing a number of beads representing the actin filaments where the persistence length is a predominant factor in the theory of diffusion coefficient as well as the electrophoretic mobility.

The approach also provides an accurate expression for the electrical polarizability, < *α* >[17].

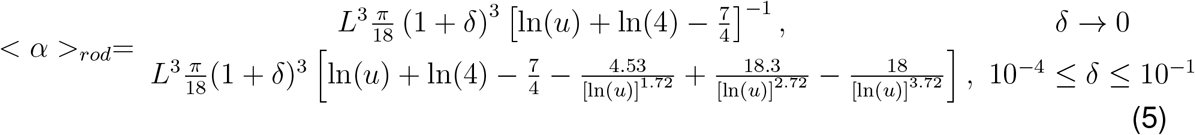

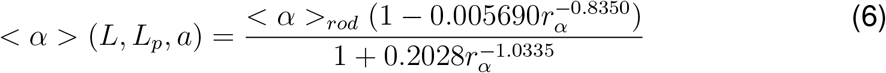

and the radius of gyration, *R_g_*

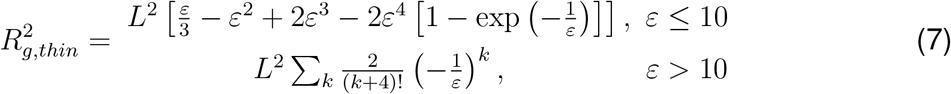

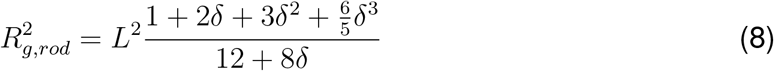

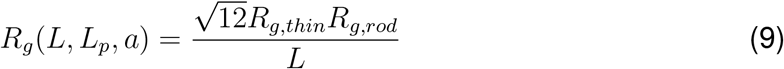

The approach generalizes previous results including Yamakawa-Fuji’s theory[20] which is accurate for long chains only. This theory was successfully tested against experimental data on double stranded DNA.

#### Electrophoretic Mobility Theory

We used Völkel’s theory[18] to calculate the monodisperse electrophoretic mobility of stiff-charged molecules in solution. In the low external electric field limit, the WLC model can be accurately represented by a semiflexible Gaussian chain consisting of *N* monomers (beads) of radius *a*, charge *q*, center-to-center monomer separation distance *b* = 4a, and persistence length *L_p_*. Whereas the aqueous electrolyte solution is considered as a homogeneous, incompressible solvent with viscosity, η, and arbitrary inverse Debye length, *κ*.

As a unique feature, this actin filament representation accounts for all the forces acting on each bead (see Figure 2) such as bending and stretching forces due to intramolecular potentials and the excluded volume and Coulomb interactions, and externally applied electric fields. Additionally, the Rotne-Prager tensor approximation[18] is used to calculate the hydrodynamic interactions between monomers, the counterion relaxation effects are neglected, and the Debye–Hückel potential[21] is used in the electrical force calculations. More, the Kirkwood-Riseman orientational pre-averaging approximation[22] is used to account for all the different rotational conformations of a filament. As a result, the expression for the monodisperse electrophoretic mobility expression for single actin filaments reads,

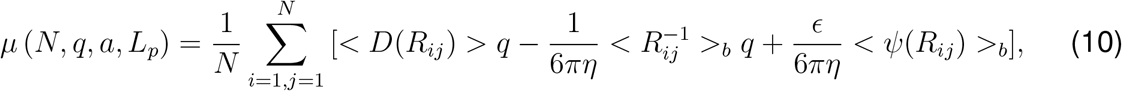

where,

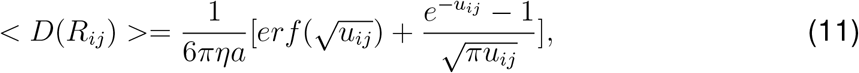

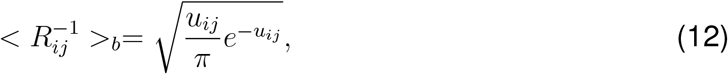

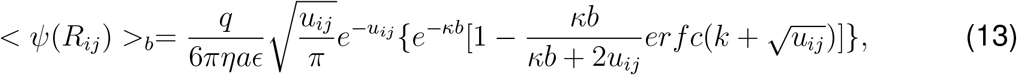

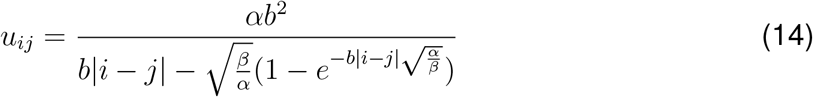

**Figure 2:**
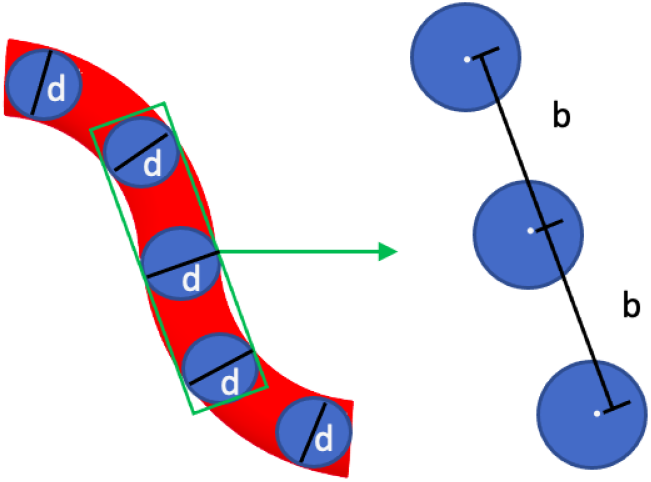
Conformation of a chain beads representing an actin filament where external forces and an electric field applied to beads form an arbitrary distribution of bending and structural conformations. The distance between beads is defined as the parameter b, as seen on the right side of the image. The chain of beads is under the effect of an electric field represented by ‘E’.

In the previous expressions, < … > represents the orientational average, 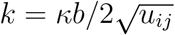, *α* = 3/4*L_p_*, *β* = 3*L_p_*/4, and *erf*(*x*) and *erfc*(*x*) are the error and complementary error functions, respectively. The first term in eq. (10) represents the monodisperse electrophoretic mobility’s contribution to the hydrodynamic interactions between monomers. The second and third terms account for the electrostatic screen generated by the electrolyte on the monomer charges.

The formulation has been validated for the single- and double-stranded DNA and numerical simulations, and it generalizes previous approaches, including the method introduced by Muthukumar[23] and Oseen[24].

#### Length Distribution Theory

The G-actin polymerization in aqueous electrolyte solutions generates filaments of different contour lengths[9, 10, 25, 26] (see Figure 3). The filament length distribution represents the number of actin filaments with a given contour length *L*. It depends on G-actin concentration, polymerization buffer, ionic strength, and the significant, independent, asymmetric, length growth rate *λ*_+_ and *λ*_−_ from both barbed and pointed ends, respectively[5, 27]. The filaments polymerize bidirectionally with the rate at the fast end about ten times larger than at the slow end. The fast end is the barbed end; the slow end is the pointed end[27]. In this work, we used the generalized Schulz distribution *Y*(*L*, *λ*_+_, *λ*_−_, *bi*) introduced by Jeune-Smith[19] for cytoskeleton filaments

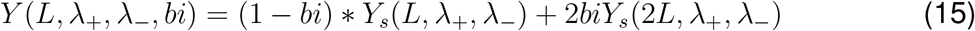

where,

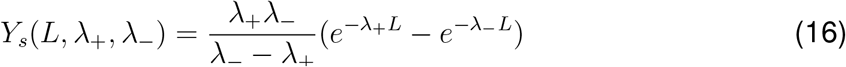

and the parameter *bi* represents the fraction of broken filaments which accounts for the shearing effects. Furthermore, we considered the experimental relationship between associate rates *λ*_+_ = 10λ_[5].

**Figure 3:**
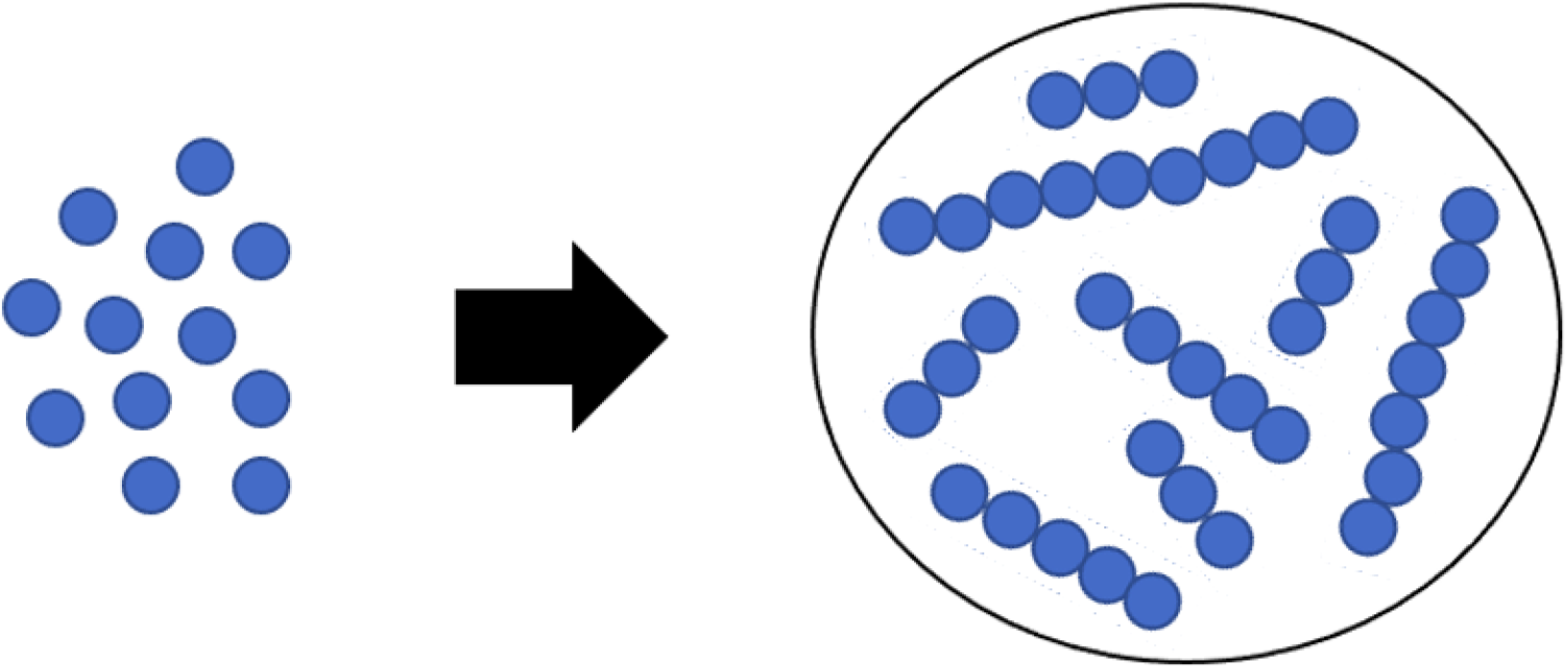
Schematic representation of G-actin monomers (left) that polymerizes into many different actin filament lengths (right). This account for the polydispersity that we encounter in a real system.

The generalized Schulz distribution was validated on microtubules polymerized in vitro, and it generalizes previous approaches, including the classic Schulz distribution theory developed for polymers with equal length distributions at each end [28]. In this work, we do not consider annealing effects on the actin filament length distribution since the sample preparation protocol used in the experimental work is designed to minimize breaking and the combination of actin filaments.

#### Polydispersity Theory

The relative contribution of individual biomolecules to some macroscopic properties, including those measured by light scattering experiments[9], is often proportional to their mass fractions *M*, in such a way that larger biomolecules gain greater significance. Considering the assumption where all actin filaments have the same diameter and density, the mass fraction of any actin filament becomes proportional to the squared contour length. Thus, we used the actin filaments weight function *M* ~ *L*^2^ and the length distribution given by eq. (16) to take the length average of eqs. (1), (10), (6), and (9). The resulting expressions for the polydisperse translational diffusion coefficient, electrophoretic mobility, gyration radius, and polarizability for actin filaments in aqueous salt solutions read

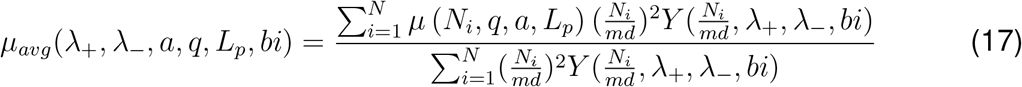

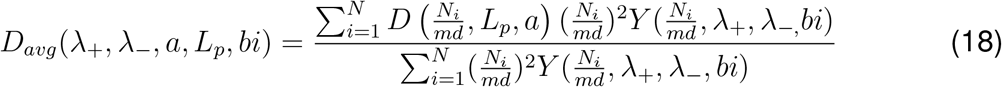

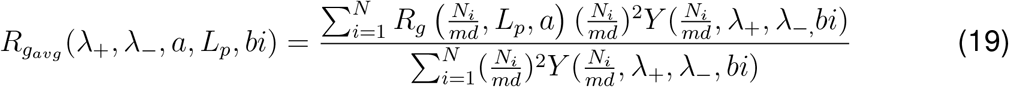

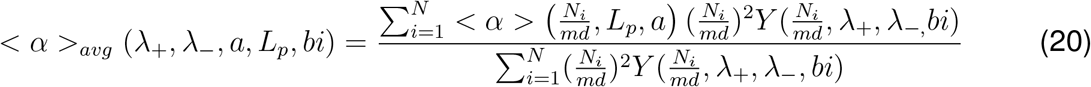

In the previous equations, we used the relationship between the degree of polymerization (the number of G-actin monomers per micrometer) *md*, the monomers number *N_i_*, and the contour length *L_i_* = *N_i_/md*. Additionally, we used the experimental value *md* = 370/*μm* [9], and generated a histogram for the filament contour length distribution using 0.2*μm* intervals (bins): 0.2*μm*, 0.4*μm*, 0.6*μm*,.., 5.4*μm*. Thus, the summation in eqs. (17) and (18) was performed over the monomers number *N_i_* = 74(*i*-1), with *i* = 2,3,.., 28.

#### Dynamic structure factor theory for semiflexible polymers

We used Kroy’s theory[11] to calculate the first cumulant (initial decay rate) *γ*_o_ and the dynamic structure factor *g_1_*(*k_s_, t*). The approach is based on the WLC model and the theory for Brownian particles in hydrodynamic solvent in dilute solutions. For short times, *g*_1_(*k_s_, t*) ~ exp [-*γ*_o_*t*] with the initial decay rate given by,

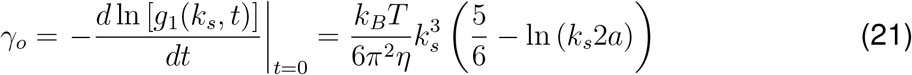

where 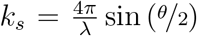 is the scattering wave number, *λ* the wavelength, *θ* the scattering angle, and *a* the filament radius. Whereas the time decay of the dynamic structure factor is given by the stretched exponential approximation

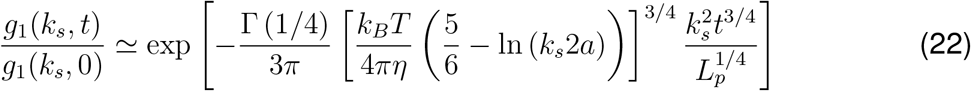

Additionally, the initial decay rate *γ*_o_ and the dynamic structure factor *g*_1_(*k, t*) can be obtained from the normalized autocorrelation function *g*_2_(*k_s_, t*) measured in DLS experiments [12, 29].

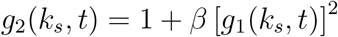

where *β* is a constant depending on the optical system used and can be determined by using the property *g*_2_(*k_s_, t* → 0) → 1.

#### Zeta Potential

We used Oshima’s approach[30], and the values for the electrophoretic mobility were measured experimentally to estimate the filament zeta potential, *ζ*. Considering actin filaments oriented at an arbitrary angle between their axis and the applied electric field, its electrophoretic mobility, *μ_avg_*, averaged over a random distribution of orientation is given by the following expression

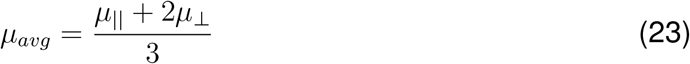

In eq. (23), *μ*_||_ represents the electrophoretic mobility for filaments oriented parallel to an applied electric field, which can be calculated using the Smoluchowski’s equation (24).

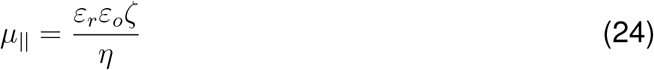

where *ε_r_* is the relative permittivity, *ε_o_* is the permittivity of a vacuum, and *η* the solvent viscosity. While *μ*_⊥_ is the electrophoretic mobility for filaments oriented perpendicular to an applied electric field. In this case, Oshima included a relaxation effect correction to Henry’s approach[31], leading to the following expression for *μ*_⊥_

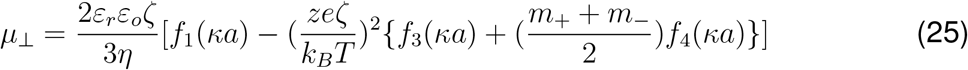

where 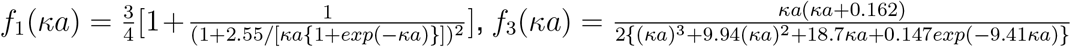, 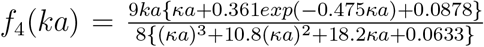. In eq. (25), *z* is the valence of counterions of the electrolyte solution, *e* is the elementary electric charge, 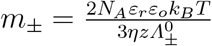 is the dimensionless ionic drag coefficient, and 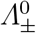 are the ionic conductance for *K*^+^ and *Cl*^-^ species.

#### Parameters calculation

The values for the set of unknown parameters *λ*_−_, *a, q, bi*, and *L_p_* usually depend on the specific electrolyte conditions, polymerization buffers, and sample preparation protocols, among other factors[8]. In this work, we found optimal values for these parameters that better reproduce the values for *γ*_0_, *g*_1_(*k_s_, t*), *μ_exp_* and *D_exp_* measured in the ELS and DLS experiments with *k_s_* = 2.6354 · 10^7^/*m* when *θ* = 173° and *λ* = 633*nm*.

In the first step, we used eq. (21) and the linear fit function for ln [*g*_1_(*k_s_*, *t*)] in Mathematica software v12.2 to determine the effective filament radius ‘*a*’. Meanwhile, substituting this parameter into eq. (22) and the use of the nonlinear fit function for

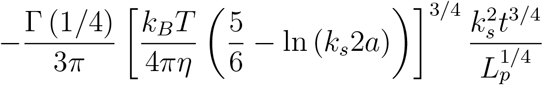

yields the value for the persistence length *L_p_* [12].

In the second step, we use Mathematica software and non-linear constrained global optimization techniques[32] to minimize the square sum cost function

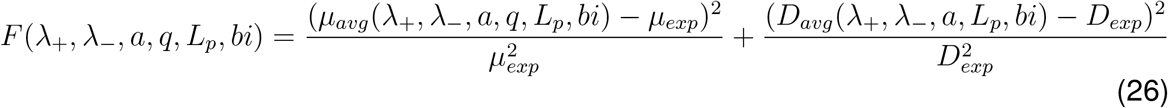

with respect to the set of parameters *λ*_, *bi*, and *q*. We found that the algorithm *“ N Minimize”* and the configuration:

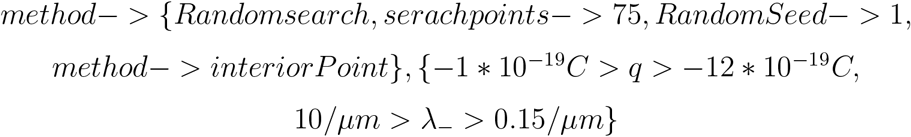

provided the most accurate and efficient minimization approach. We used the random search algorithm, which generates a population of random starting points and uses a local optimization method to converge to a local minimum. Then, the best local minimum is chosen to be the solution. We used 5, 10, 25, 50 and larger numbers of search points. We found that numbers of search points larger than 75 generated the same optimal values. Further, we used the nonlinear interior point method, one of the most powerful algorithms to find the local minimum of a sum of squares[33]. Additionally, we used the numbers (0, 1, 5, 10) for the RamdonSeed parameter to consider different starting values in the random number generator algorithm. We found that the optimal values usually did not depend on these numbers. We constrained a range in the values of the parameters to avoid those with unphysical meaning and bracketed those typical values found in the literature. We used the *“ParallelSum”* and *“RemoteKernel”* Mathematica functions to run the Mathematica notebook on a computer cluster with 44 cores and 140Gb RAM.

## II. EXPERIMENTAL SECTION

### A. Sample preparation

Actin from rabbit skeletal muscle (> 99% pure) was purchased from Cytoskeleton Inc. and used without further purification. We prepared three actin filament samples using the G-actin buffers, polymerization buffers, and electrolyte solutions tabulated in tables I, II, and III, respectively. We used the same sample preparation protocol for each sample. A 1.0 *mg* of actin powder was reconstituted to 10 *mg/ml* G-actin density by adding 100 *μL* of Ultra-pure Distilled water Molecular Biology. Next, we added 2.40*mL* of G-actin buffers (see table I), aliquoted into experimental samples, and stored in cryo-tube vials at −70° *C*. The G-actin solutions were incubated on ice for one hour to de-polymerize actin oligomers that may be formed during storage before polymerization. 20 *μL* of polymerization buffers (see table II) were added to 200 *μL* G-actin solutions and transferred into Beckman coulter centrifuge tubes for one hour at room temperature to finish the polymerization stage. By balancing the needs of sample preservation and rapid run time, we centrifuge each experimental sample for two hours using the Allegra 64R Benchtop Centrifuge (Beckham Coulter) at 4°*C* using a speed of 50,000 G-force. Following this process, 22 *μL* of protein pellet was obtained by extracting 198 *μL* of the un-wanted supernatant (90%). Consequently, we added 978.0*μL* of electrolyte solution (see table III) to the pellet leading to a final volume of 1.0mL, and stored the final solution at 4° *C* overnight to achieve polymerization equilibrium in our samples. The pipetting tips used in our experiments were cut to an average diameter of ~ 5–7*mm* and prevent filament breakage[3, 9]. The pH of G-actin buffer, polymerization buffer, and electrolyte solutions were adjusted by adding either Hydrochloric acid (HCl) volumetric standard or Sodium Hydroxide (NaOH), pellets 97+%, A.C.S. reagents from Sigma Aldrich. The pH was measured with an accuracy of ±0.002 using Thermo scientific Orion Star™ A211 Benchtop pH Meter. We also determined the actin protein concentration experimentally using spectrophotometer techniques[15] and the Precision Red Advanced Protein Assay Reagent (Cat# ADV02)[34, 35] from Cytoskeleton.inc. We obtained a protein concentration of 1.32*μM* across all our experiments.

**Table I:** Table for G-actin buffers

|  | Buffer 1 | Buffer 2 | Buffer 3 |
| --- | --- | --- | --- |
| Tris Base | 2 mM | 2 mM | 2 mM |
| CaCl<sub>2</sub> | 0.2 mM | 0.2 mM | 0.2 mM |
| BME (Beta-Mercaptoethanol) | - | 0.1 mM | - |
| ATP (Adenosine triphosphate) | 0.5 mM | 0.2 mM | 0.5 mM |
| DTT (Dithiothreitol) | 0.2 mM | - | 0.5 mM |
| pH | 7.80 | 7.66 | 8.23 |
| Ionic Strength | 0.0057 | 0.0037 | 0.006 |

**Table II:** Table for Polymerization Buffers

|  | Buffer 1 | Buffer 2 | Buffer 3 |
| --- | --- | --- | --- |
| KCl | 150 mM | 150 mM | 50 mM |
| MgCl<sub>2</sub> | 2 mM | 2 mM | 2 mM |
| pH | 7.56 | 7.64 | 8.07 |
| Ionic strength | 0.155 | 0.155 | 0.055 |

**Table III:** Table for Electrolytes

|  | Buffer 1 | Buffer 2 | Buffer 3 |
| --- | --- | --- | --- |
| KCl | 0.1 M | 0.1 M | 0.1 M |
| pH | 7.72 | 7.66 | 8.06 |
| Ionic strength | 0.1 | 0.1 | 0.1 |

### B. Light Scattering Experiments

We used Malvern ULTRA Zetasizer instrument equipped with a He-Ne 633 nm laser to measure actin filament’s translational diffusion coefficient and electrophoretic mobility at low G-actin protein concentrations. The experiments were configured, and the data was recorded and analyzed using Zetasizer Xplorer software. The ULTRA Zetasizer features an Adaptive Correlation algorithm that uses information from the sample to determine how long it measures to ensure data consistency. This feature also applies intelligent logic to separate erroneous data associated with transient artifacts such as dust or aggregates. Adaptive Correlation intelligently identifies rogue large particles and filters these from the presented data but retains consistently present populations. In all our measurements, we used 180 seconds as the equilibration time to thermally stabilize the sample at the desired temperature of 25° *C*. The Zetasizer instrument uses a cell compartment that keeps the temperature constant during the scattering measurements. Additionally, the attenuation factor was set to automatic using 11 positions to control the beam intensity from 100% to 0.0003%. In this mode, the instrument showed an attenuation factor between 10 and 11 across all measurements during our DLS and ELS experiments. We also selected the *“protein”* material option with a refractive index of 1.450 and absorption of 0.001. Furthermore, we selected *“water”* as the dispersant option with a refractive index of 1.33, and a viscosity of 0.8872*mPa.s*. It is worth mentioning that the refractive index and absorption of the material have no bearing on the Z-average, polydispersity, and intensity distribution results.

In the DLS experiments, 1.0 *ml* of actin filament solution was collected in the 12 mm square Polystyrene cuvette (DTS0012). The correlation functions were measured at the Back-scattering angle (173°), where the incident beam does not have to travel through the entire sample, and the effect of multiple scattering and dust is greatly reduced. We ran five consecutive, independent experiments for each actin filament sample to reduce statistical errors in the translational diffusion coefficient values. We calculated the average of the three longitudinal diffusion coefficient values with the lowest standard deviation and better correlation function to minimize error and increase reproducibility. The measurement duration was automatically determined from the detected count rate. In this mode, the lower the count rate, the longer the measurement duration, and the higher the noise. We used the ‘General Purpose’ analysis model, which uses a non-negative least squares (NNLS) analysis. It is the more suitable model for our case due to the unknown size distribution.

In the ELS experiments, we used the Malvern Panalytical Universal dip cell kit (ZEN1002) and the 12 mm square Polystyrene cuvette (DTS0012) for measuring the electrophoretic mobility of actin filaments. We ran three independent experiments for each actin filament sample to reduce statistical errors in electrophoretic mobility values. The voltage selection and measurement process were set to automatic. The Zetasizer Xplorer software automatically measures the sample electrical conductivity in this mode. It adjusts the cell voltage to keep a low current flowing, close to 5 mS/cm, in the sample. Otherwise, the sample temperature may increase near the electrodes, inducing bubble formation, sample degradation and, consequently, misleading data measurements. The software automatically selects the most appropriate analysis and collection data model based on the cell type chosen, dispersant properties, and the sample’s conductivity. We focused on the fast field reversal (FFR) of the phase analysis light scattering (see figure 6) since the mobility measured during this period is due to the electrophoresis of the particles only. It is not affected by electro-osmosis associated with the soft field reversal (SFR).

**Figure 4:**
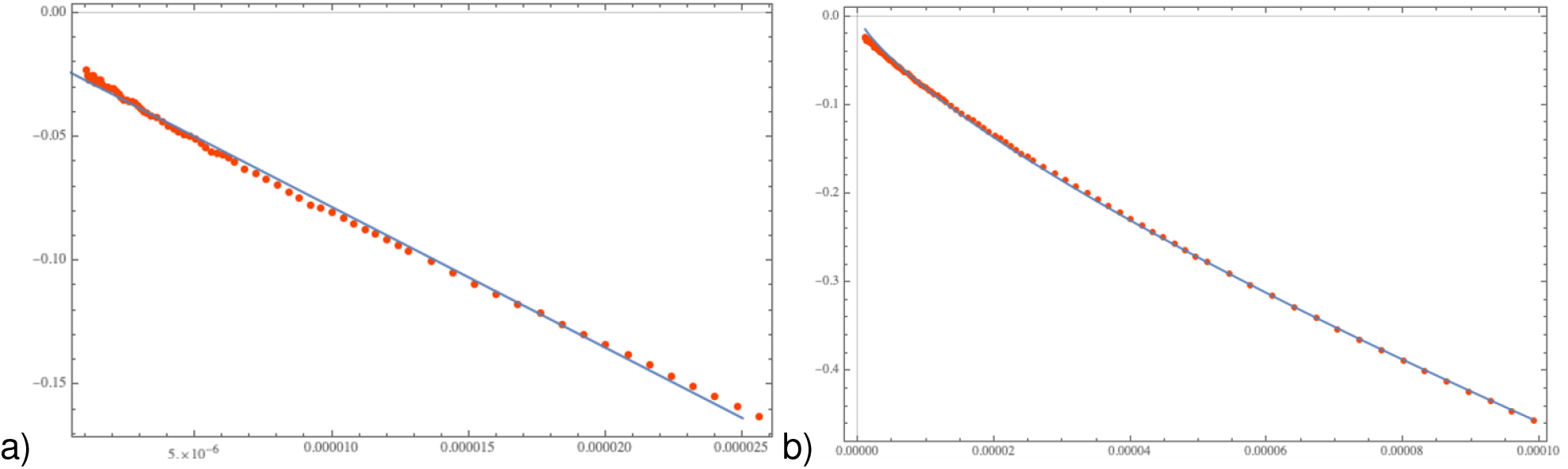
Mathematic Fitting Approach: a) Initial decay rate (*γ*_0_), and b) Persistence length (*L_p_*) fitting Mathematica plots using eqs (21) and (22).

**Figure 5:**
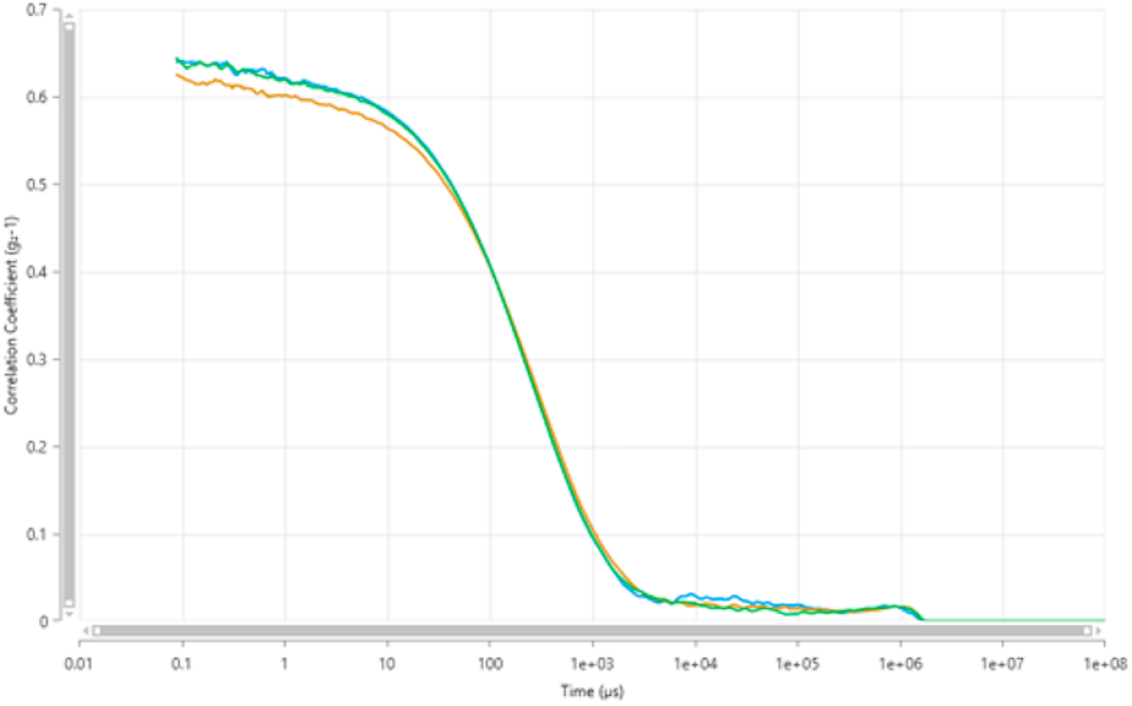
Correlation plot. Z-average of 190 nm ± 8.3 nm, polydispersity index (PDI) of 0.56, derived mean count rate 366 ± 15.7 kpcs, and a diffusion coefficient of 2.60 ± 0.111 *μm*^2^/*s*.

**Figure 6:**
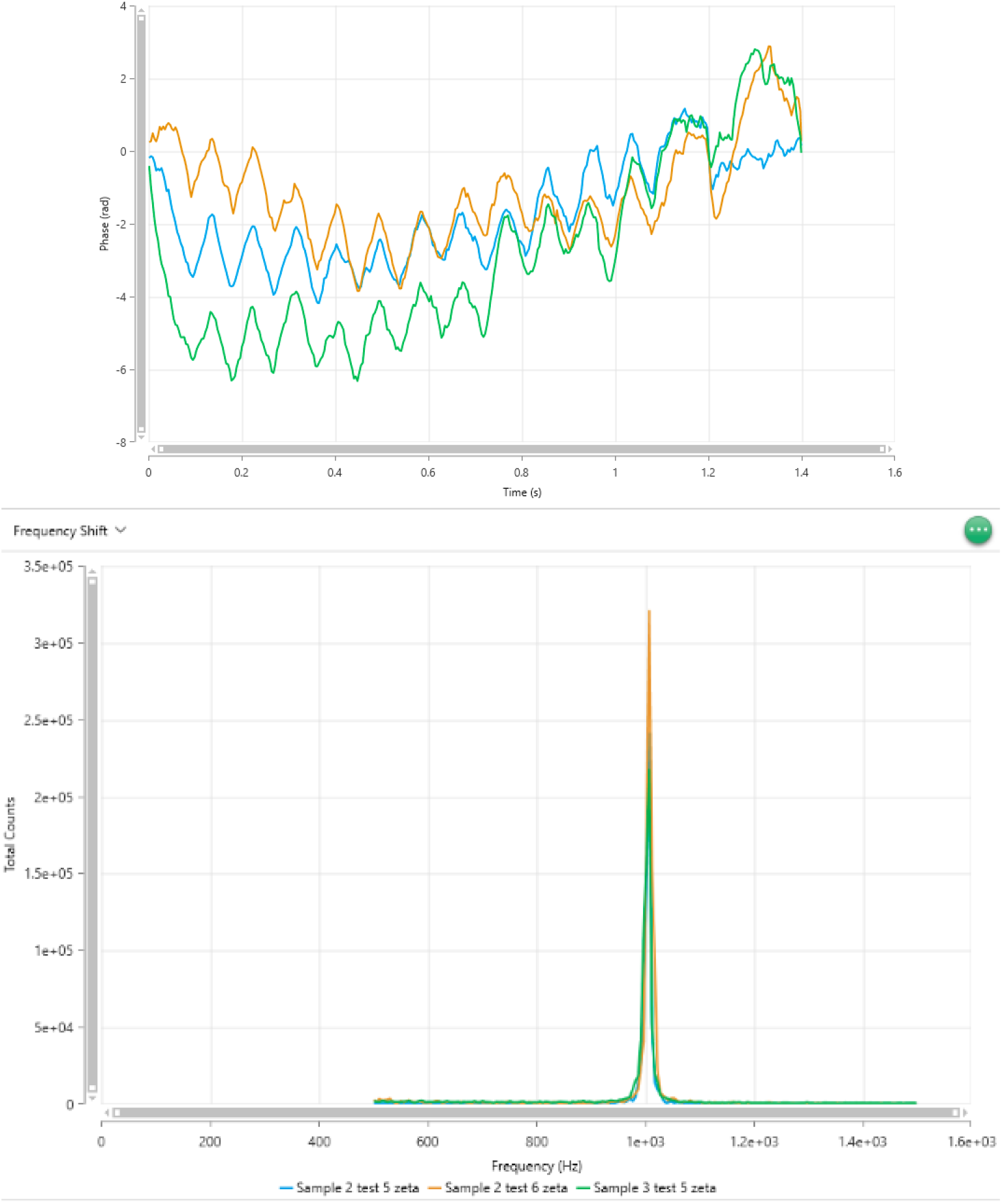
phase plot and frequency plot for buffer 3 with quality factor 1.37± 0.321, zeta potential (mV) of −13 ± 0.506, conductivity (mS cm-1) of 12.4 ± 1.69, mobility (μm·cm / V·s) of −1.02 ± 0.0395.

We improved reproducibility by including a pause between consecutive measurements. A time delay also helped reduce sample heating, allowing the sample to recover 25°C between consecutive measurements, reducing critical sample’s degradation, and avoiding increasing mobility with sequential measurements. The minimum and maximum repeat runs per experiment were manually set to 10 and 30, while the pause duration and pause between repeats were set to 10 and 60 seconds, respectively.

## III. EXPERIMENTAL RESULTS

The translational diffusion coefficients and polydispersity index (PDI) are obtained from the correlation function of the scattered intensity. An illustrative plot is depicted in figure (5). The average diffusion coefficient values with average standard deviation and percent error are summarized in table (IV). The diffusion coefficients of actin filaments were obtained at different pH, reducing agents, ionic strength, and ATP concentration. All correlation plot results and intercepts were lower than one and within a range in the low polydispersity index of 0.3 - 0.5, indicating a good quality in our samples. Additionally, all our experimental size distribution measurements displayed a derived count rate higher than 100 kpcs, the minimum value required to obtain suitable measurements.

**Table IV:**
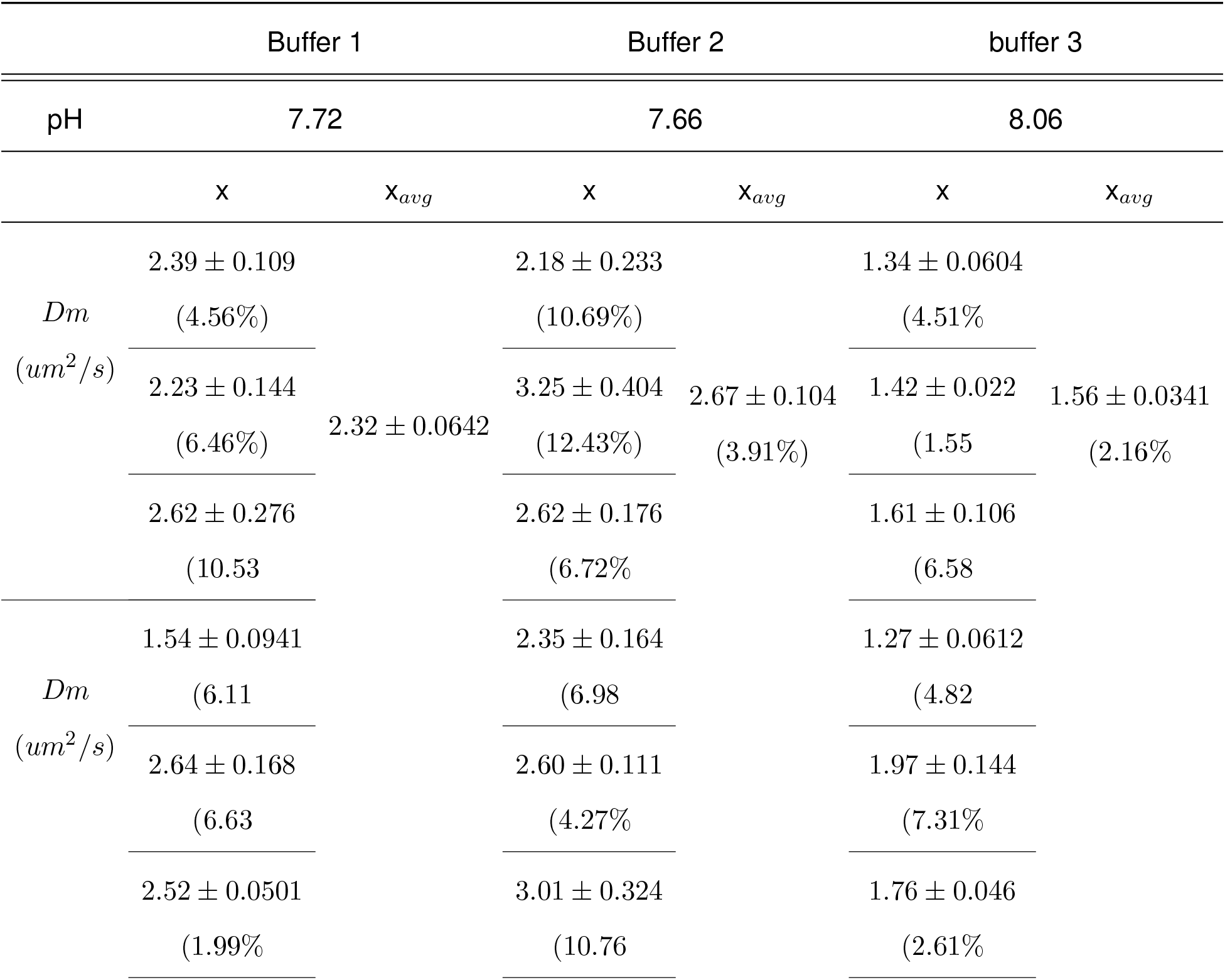
Experimental results for Diffusion coefficients

The values obtained for the average electrophoretic mobility with standard deviation and percent error are tabulated in table (V). In figure (6), we show an illustrative example on the graph obtained for the electrophoretic mobility using buffer #3.

**Table V:** Experimental results for Electrophoretic mobility

|  | Buffer 1 |  | Buffer 2 |  | Buffer 3 |  |
| --- | --- | --- | --- | --- | --- | --- |
| pH | 7.72 |  | 7.66 |  | 8.06 |  |
|  | x | x<sub>avg</sub> | x | x<sub>avg</sub> | x | x<sub>avg</sub> |
| Electrophoretic Mobility (um.cm/V.s) | -0.941 | -0.955 ± 0.014 | -1.04 | -0.955 ± 0.0775 | -0.986 | -1.02 ± 0.0378 |
|  | -0.969 | (1.47% | -0.938 | (8.12% | -1.01 | (3.70% |
|  | -0.955 |  | -0.888 |  | -1.06 |  |
| Zeta Potential (mV) | -14.150±0.206(1.46%) |  | -14.082±1.137(8.07%) |  | -14.265±0.527(3.69%) |  |

The quality factor is a parameter that derives from the phase analysis during the FFR stage of the measurement. All our experimental electrophoretic mobility results were obtained with a quality factor in the range of 1.08 - 1.37. These values are more significant than 1, which is the minimum value required to obtain good data quality. Another evidence of good data quality is displayed in our frequency shift plots since there are no traces of noise, and the plots match very well.

Using the correlation plot data of six independent sets of three runs each per buffer (see figure 4), and the fitting approach described in the section “Parameter calculation,” we obtained the following results for the initial rate decay, the filament radius, and the persistence length *L_p_* for each buffer solution

## IV. THEORETICAL RESULTS

The optimal values for the parameters *λ*_−_, *b* and *q* are tabulated in table VII. The values for *λ*_−_, *λ*_+_ and *b* were used to calculate the weight length 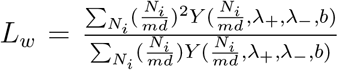 and number length 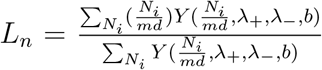 averages. We also calculated the polydispersity number *PDI* = *L_n_/L_w_*, the average hydrodynamic radius 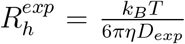, and the length distribution *Y*(*L, λ_+_, λ_−_, b*). The values for the parameters *λ_+_*, 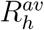, *PDI, L_n_*, and *L_w_* are given in table VIII.

**Table VI:** Calculations of *d, L_p_*, and *γ*_0_ from equations 21 and 22

| *Buffer* | $\gamma_0[1/s]$ | $d[nm]$ | $L_p[um]$ |
| --- | --- | --- | --- |
| 1 | $6056.74 \pm 46.31$ | $10.77 \pm 0.168$ | $0.632 \pm 0.00376$ |
| 2 | $5883.99 \pm 66.99$ | $11.27 \pm 0.263$ | $0.633 \pm 0.00797$ |
| 3 | $3479.51 \pm 33.98$ | $26.29 \pm 0.306$ | $0.817 \pm 0.00676$ |

**Table VII:** Optimal parameters

|  | Buffer # 1 | Buffer # 2 | Buffer # 3 |
| --- | --- | --- | --- |
| $\lambda_-[1/\mu m]$ | 1.734 | 1.890 | 1.361 |
| $1/\lambda_-[\mu m]$ | 0.5966 | 0.5291 | 0.735 |
| $a[nm]$ | 5.385 | 5.635 | 13.145 |
| $q[e]$ | -4.748 | -4.974 | -12.138 |
| $L_p[\mu m]$ | 0.632 | 0.633 | 0.817 |
| $b$ | 0.481 | 0.685 | 0.467 |

**Table VIII:** Calculations using the optimal parameters

|  | Buffer # 1 | Buffer # 2 | Buffer # 3 |
| --- | --- | --- | --- |
| $\lambda_+[1/\mu m]$ | 17.34 | 18.90 | 13.61 |
| $R_h^{av}[\mu m]$ | 0.106 | 0.0923 | 0.157 |
| $PDI$ | 0.568 | 0.585 | 0.551 |
| $L_n[\mu m]$ | 0.562 | 0.472 | 0.687 |
| $L_w[\mu m]$ | 0.989 | 0.807 | 1.248 |

Whereas, the filament length distributions *Y*(*L, λ_+_, λ_−_, b*) are shown in Figure 8.

**Figure 7:**
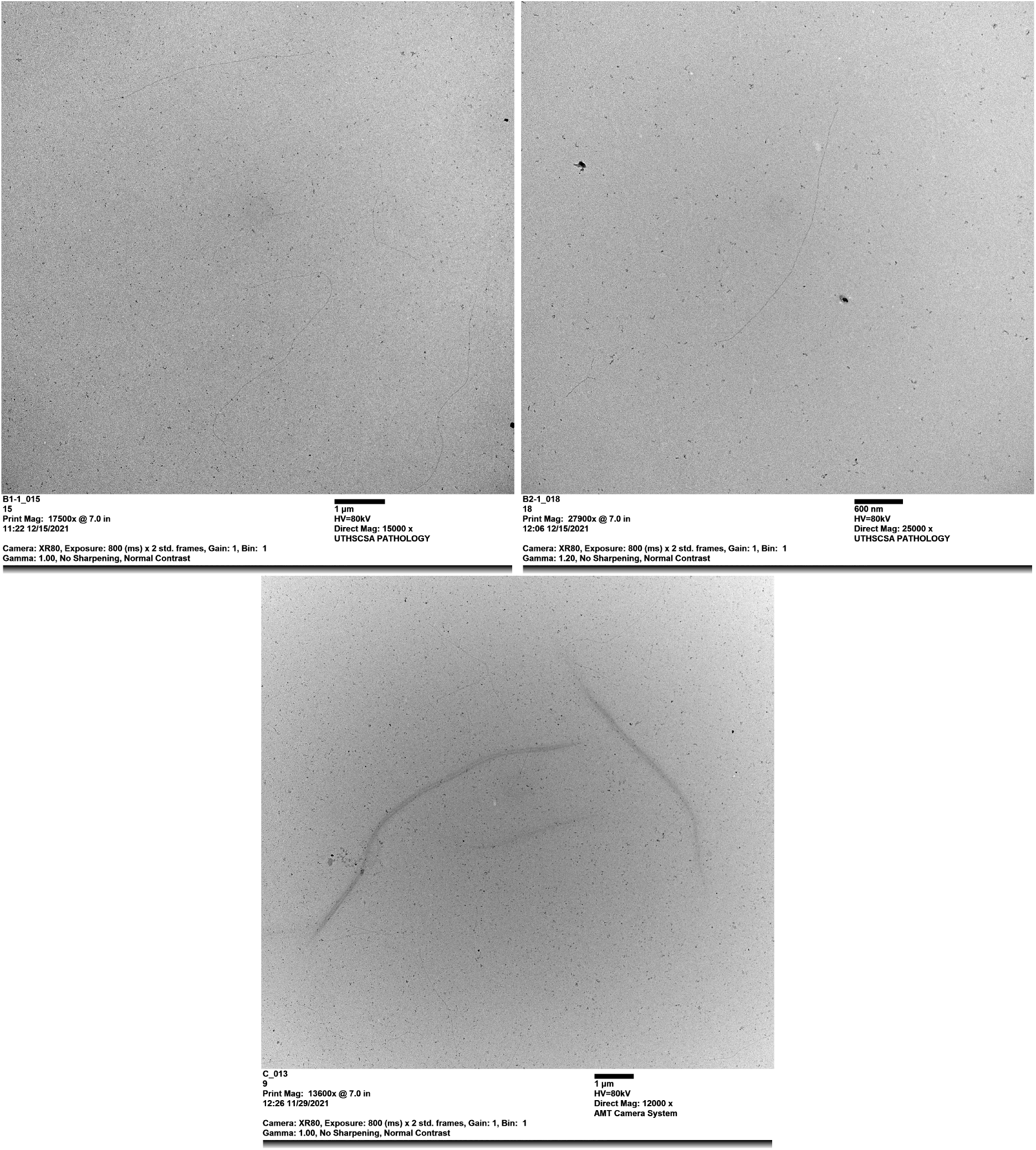
Micrograh Images for buffer #1, #2, and #3. These images were taken from the JEOL 1400 TEM.

**Figure 8:**
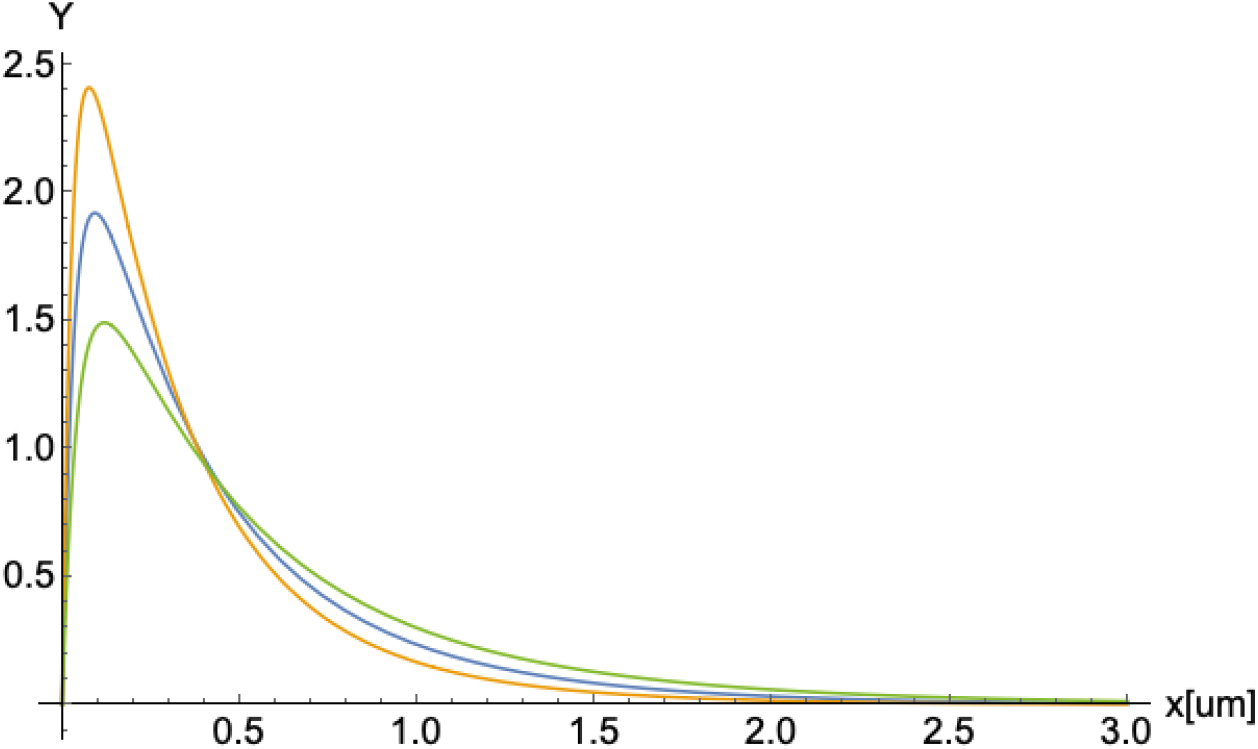
Filament length distribution *Y*(*λ_+_*, *λ*_−_, *L*). Blue, orange, and green colors represent buffers #1, #2, and #3 respectively.

## V. DISCUSSION

### A. Length Distribution

Our results for the weight- and the number-average length *L_w_* and *L_n_* varied for all cases; however, the polydispersity index (PDI) ratio remains constant among the three buffers, which agrees with previous results[9, 25]. Based on the structure and mobility of actin filaments, Janmey’s group[9] observed a formation of long filaments in the F-actin’s length distribution as they increased the actin/gelsolin molar ratios. Burlacu’s group[25], used electron micrographs to analyze the length distribution of actin filaments under the presence of phalloidin-A, and IATR-actin. Both research groups obtained high length rate values, but the PDI remained the same for all cases. We noticed that the increase in the polymerization-growth rate of filaments for buffer #1 led to an increase on *L_n_* and *Lw* compared to the results obtained for buffer #2. These differences may be, in part, due to the chemical composition of the two buffers. One main difference between these two buffers is the presence of 1.4-Dithiothreitol (DTT) and Beta-Mercaptoethanol (BME) concentrations. To illustrate, buffer #1 has 0.2 mM DTT reducing agent, whereas buffer# 2 contains 0.1 mM BME. These two reducing agents are essential to prevent the formation of oligomers and agglomeration of monomers and maximize the availability of free G-actin monomers for polymerization[36]. Further, DTT is more efficient than BME in lowering F-actin’s storage modulus, e.g., the overall resistance to deformation. Thus, the DTT concentration used in buffer #1 could partially compensate and generate effects similar to the BME concentration used in buffer #2. Another critical difference is that buffer #1 has 2.5 times higher ATP concentration than buffer #2. While the actin addition (elongation) rate depends on free ATP-G-actin concentration, the subunit loss rate does not[5, 37], meaning that buffer #1’s ATP monomer pool is more significant than #2. Although buffer #1’s growth association rates are higher than buffer #2’s by 11%, the polydispersity index (PDI) values remain similar for buffer #1 and #2. Additionally, the shearing parameter, *b*, associated with the breakage fractioning of actin filaments in solution, is higher for buffer #1 than buffer #2 (see figure 8). We correlate this result to an increase in actin filament lengths. Indeed, the shearing effects is somewhat proportional to the filament lengths, where the more prominent the filaments grow, the more filaments are exposed to shear and break.

Interestingly, buffer #3 revealed the formation of much longer filaments compared to the other buffers caused by an increase in the association rates (see table VII). Buffer #3’s association rates differ from buffer #1 and #2 by 18.83% and 28.01%, respectively. An essential difference is the 2.5 times higher DTT concentration of buffer #3 than buffer #1. We correlate the DTT’s efficiency and increment in concentration to an increase in actin filament’s length formation, leading to an increase in elongation rates. In addition, the increase in ATP concentration leads to an increment of free ATP-G-Actin free monomers in the solution. Thus, the *L_n_* and *L_w_* parameters in buffer #3 have higher values than the other buffers (see figure VII). Unlike buffer #2, the shearing parameter for buffer #3 resembles buffer #1’s result, where the filaments are more commonly fractioned due to the longer average filament lengths. Therefore, we conclude an inverse proportionality correlation between the shearing parameter and the DTT concentration in the solution.

### B. Structural Parameters

We evaluated 18 experimental autocorrelation plots (see figure 5) obtained from dynamic light scattering measurements using the dynamic structure factor theory. For each buffer, we extracted the initial decay rate, γ_0_, in a time frame ranging ~ 10^-6^ to 3 × 10^-5^ seconds and used eq. (21) to obtain the effective diameter, *d*. The corresponding values are shown in tables VI and VII. They agree with previous work in hydrodynamic conditions[12, 38]. However, they are more significant than those obtained from bare molecular structure filament models. The use of Cong molecular structure model[39] for 13 polymerized G-actin monomers (see figure 9a) and the approach for an effective cylindrical model described by Marucho’s group[33] yield an average filament diameter of *d_MS_* = 4.77*nm*. As a result, the difference between the effective and bare diameters is equal to Δ*d* = 10.77*nm* – 4.77*nm* = 6.0*nm* and Δ*d* = 11.27*nm* – 4.77*nm* = 6.5*nm* for buffers #1 and #2, respectively. The increase in diameter can be explained using the MacMillian-Mayer theory for highly charged colloidal cylinders in monovalent salt solutions[40]. The approach predicts that the effective filament diameter is equal to the summation of the bare diameter, *d_MS_*, and the contribution, Δ*d*, from the filament charge and the electrical double layer (EDL) surrounding its surface. In particular, the calculation for a rod-like cylinder with uniform linear charge density *λ* = −4*e/nm*, and diameter *d_MS_* = 4.77*nm* immersed in 0.1M monovalent salt solution (KCl) yields an increment in the size of *Δd* = 5.49*nm*, which is similar to the values obtained in our previous calculations. The effective (integrated) monomer charges presented in the table VII are smaller than those charges (−12e) obtained from bare G-actin molecular structures. This is due to the charge attenuation coming from the electrostatic screen generated by the high accumulation of counterions around the filament surface[33, 41–43].

**Figure 9:**
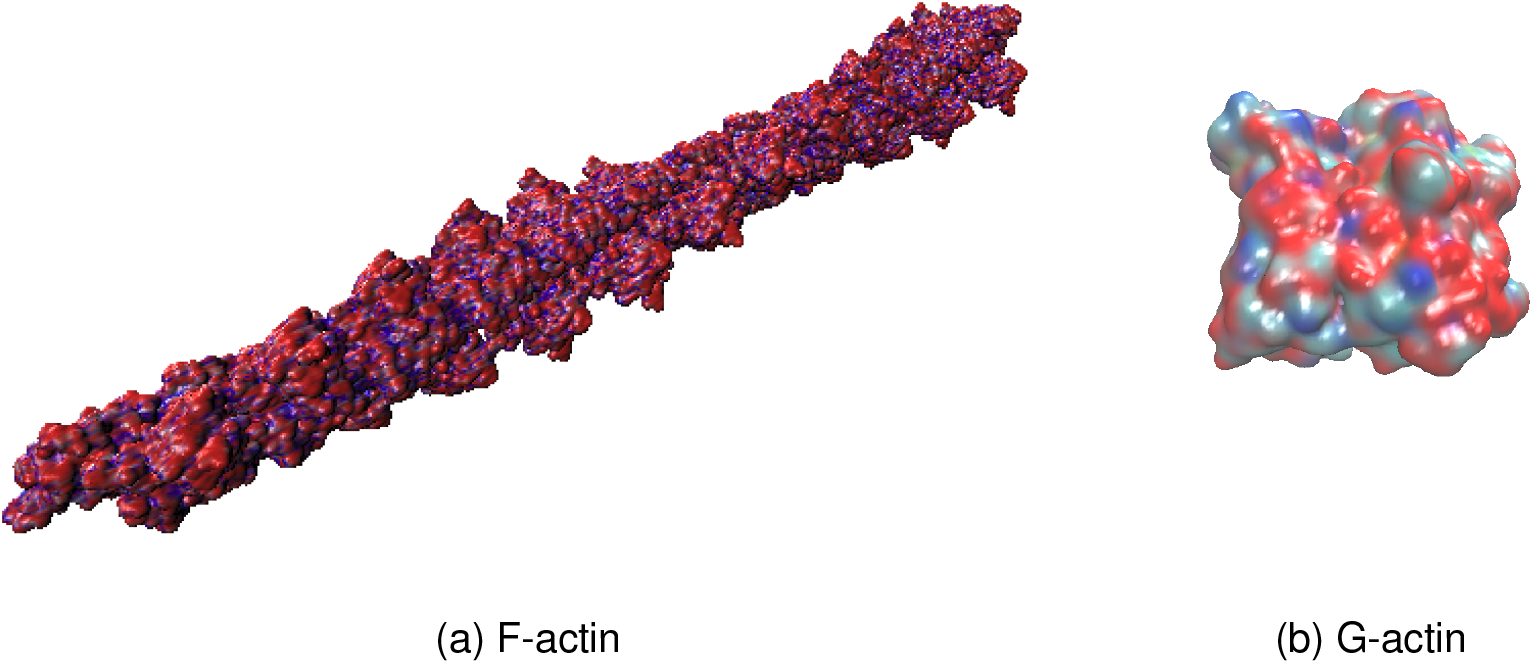
Molecular structure models

Interestingly, the effective diameter of 26.29 *nm* obtained for buffer #3 is more significant than for buffers #1 and #2. We performed Transmission Electron Microscopy (TEM) experiments to obtain micrographs images on the three buffers (see figure 7). We observed the formation of single filaments for buffers #1 and #2; nevertheless, buffer #3 shows the formation of single filaments and the formation of actin filament bundles of different diameters. The combination of high ATP concentration, high DTT concentration, and low KCl concentration in polymerization buffer #3 could be the most impactful contributing factors in forming actin bundles. According to Lior Havir[44], an actin bundle’s diameter has a minimum of three times thicker than a single filament’s diameter, which is in agreement with our results for buffer #3. Additionally, the formation of longer filaments could lead to actin bundles in solution [26]. This assumption fits well with our results since buffer #3 produces longer actin filaments leading to smaller diffusion coefficients than buffers #1 and #2. Additionally, Tang’s group [26] used different KCl concentrations of 30 mM, 50 mM, 100 mM, and 150 mM to induce actin bundles. Their findings show that actin bundles may form more efficiently at low concentrations of KCl. These findings agree with our results, since buffers #1 and #2 have 150 mM KCl concentration, whereas buffer #3 has 50 mM KCl only.

Finally, we extracted the values for the persistence lengths, *L_p_* using eq. (22) and the values obtained for the effective diameters. The persistence length obtained for Buffers #1 and #2 s are 0.632*μm* and 0.633*μm*, respectively. These values are in good agreement with previous experimental work[12, 19, 45]. However, buffer #3’s persistence length increments by ~ 22.64% from the previous values. This increase is due to the higher presence of free ATP-G-actin monomers in the solution. Compared to reported electron micrographs data, the lower values for the persistence length and average contour filament length obtained in this work arise from the significant difference in the association rates at the filament ends that shift to sub-micro lengths, the maximum of the length distribution. In contrast, the exponential decay of the tail of the length distribution can only be measured experimentally due to microscopy resolution limitations [19].

### C. Translational Diffusion Coefficient and Electrophoretic Mobility

Quasielastic light scattering (QLS) experiments were performed by Janmey’s group[9] to measure the translational diffusion coefficient of actin filaments using similar chemical compositions for the g-actin and polymerization buffers leading to diffusion coefficient results that agree with our results (see table IV). However, these values can significantly increase or decrease when considering different experimental protocols, chemical compounds, storage, and preparation of actin monomers, polymerization, and techniques such as fluorescence photobleaching recovery (FPR), pyrene-labeled fluorescence, fluorescence and video microscopy[1–4, 6]. For instance, Wang’s diffusion coefficient results differ from ours by 2-3 orders of magnitude because they measured the diffusion coefficient before the polymerization equilibrium was reached. In the same way, Kas’s group[3] analyzed the diffusion coefficient through the tube model[46] and the concept of reptation[47], where a tube is embodied around a single filament. They used a much higher concentration of ATP and a concentration of actin up to three times larger than ours, generating longer filaments of about ~ 20 – 50μm in length. As a result, they obtained an arithmetic mean of the diffusion coefficient, which is two orders of magnitude lower than our experimental results.

While our results for buffers #1 and #2 are pretty similar, buffer #3 showed a lower translational diffusion coefficient value of 1.56*μm*^2^/*s*, suggesting that actin filaments are longer in average. This is due to the high concentration of ATP and dithiothreitol (DTT), leading to more free ATP-G-actin monomers in the pool. Consequently, buffer #3 leads to an increase in the associated growth rate of filaments. As the filaments increase in length, the translational diffusion coefficient decreases, according to Zimmerle’s results[48].

On the other hand, Takatsuki and Li[49, 50] have performed electrophoresis experiments using actin-labeled fluorescent dyes. Since they used buffers similar to ours, they have obtained an electrophoretic mobility value of −0.85 ± 0.07*μm.cm/V.s*, which agrees with our experimental results (see table V). Following Oshima’s approach[30], we also predicted the zeta potential (ZP) from the experimental electrophoretic values for each buffer. High ZP (≳ 0.25*V*) has been commonly associated with highly charged particles inducing intermolecular repulsion and leading to dispersion stability[51]. In contrary, low ZP will likely lead to the aggregation of charged monomers. The predicted zeta potential values are similar among all buffers; however, these findings could not explain the formation of bundles in buffer #3 since other factors must be considered.

Overall, while our results for the longitudinal diffusion coefficient mainly depended on the length distribution, effective diameter, semiflexibility, chemical compounds, and reducing agents comprising G-actin buffers, the electrophoretic mobility was predominantly affected by the effective filament charge, the pH level, and the ionic strength.

### D. Other Properties

According to Steinmetz’s group[52], a comparison of Ca-G-actin, EGTA-G-actin, and Mg-G–actin polymerized with 100 mM KCl was studied to establish the impact of phalloidin over actin in a 2:1 molar concentration. The experimental protocols and buffers differ from ours by the sole presence of imidazole, NaN3, and EGTA. Similarly, De La Cruz’s research is based on the structure of nucleotide-free actin filaments [53]. They found a radius of gyration around ~ 2.4 – 2.5 *nm* in the presence of phalloidin, which is the same order of magnitude as our results (see table IX).

**Table IX:** Parameters were calculated from equations (6) and (9).

|  | Buffer # 1 | Buffer # 2 | Buffer # 3 |
| --- | --- | --- | --- |
| Radius of Gyration, $R_g[\mu m]$ | 0.348 | 0.301 | 0.440 |
| $R_{g,rod}[\mu m]$ | 0.451 | 0.378 | 0.563 |
| Polarizability, $\langle \alpha \rangle^{1/3}[\mu m]$ | 0.558 | 0.500 | 0.734 |
| $\langle \alpha \rangle_{rod}^{1/3}[\mu m]$ | 0.702 | 0.618 | 0.904 |

We calculated the radius of gyration and the polarizability, which depended on many factors such as the association rates, effective diameter, length distribution, persistence length, and the shearing effects. The radius of gyration for buffer #1 is 13.51% higher than in buffer#2 due to the increase in association rates associated with an increment of free ATP-G-actin monomers in the system, and consequently, an increase in the length distribution. Moreover, the fractioning of filaments between these buffers is ~ 30%, leading to a contributing factor in this difference. In the same way, association rates, length distribution, and effective diameter in buffer #3 are greater than in the other two buffers since we obtained longer filaments and the formation of actin filament bundles in the system. Consequently, these parameters increased 20.91% and 31.59% compared to those in buffers #1 and #2. Similarly, we analyzed the polarizability parameter of buffer #3 being 23.98% and 31.88% higher than those in buffers #1 and #2, respectively. This is due to the impact on polarizability of the intermolecular dispersion forces and the electron cloud distortion under the presence of an electric field[54]. As the filament becomes more elongated, more charges/electrons are easily moved within the e-cloud/layers, increasing their polarizability and strengthening the dispersion forces, unlike compact molecules where all charges are symmetrically together. As a result, the formation of longer filaments generates higher *<α>* and *R_g_* values.

We also analyzed the *R_g,rod_* and < *α* >_*rod*_ parameters (see table IX) for the rigid-rod case to understand the relevance of persistence length, *L_p_* in our calculations. When the persistence length is disregarded, buffer #1’s rod values, *R_g,rod_* and < *α* >_*rod*_, are 22.83% and 20.51% higher, respectively. Additionally, buffer #2 and #3’s parameters *R_g_* and *<α>* decrease 18 – 21% from those corresponding to rod values when considering their persistence length values. We concluded that actin filament semiflexibility contributes ~ 20% in the value of these parameters.

## CONCLUSIONS

In this article, we introduced a unique approach that combines light scattering experiments and optimized theoretical approaches to characterize actin filaments’ polyelectrolyte and hydrodynamic properties. We used Malvern ULTRA Zetasizer instrument to measure actin filament’s translational diffusion coefficient and electrophoretic mobility at low protein concentration. We developed a novel sample preparation protocol based on bio-statistical tools to minimize errors and assure reproducibility in our results. We also considered three different buffers, g-actin and polymerization, used in previous works, to elucidate the impact of their chemical composition, reducing agents, pH values, and ionic strengths on the filament properties.

Additionally, we optimized a novel multi-scale approach to calculate hydrodynamic and polyelectrolyte properties of polydisperse actin filaments in aqueous salt solutions. Most conventional approaches for biopolymers solutions center on rigid, monodisperse, and sometimes uncharged cylindrical models and theories. These approaches may be inappropriate for cytoskeleton filaments because they omit essential hydrodynamic and polyelectrolyte filament properties. In this article, we extended those approaches to account for filament polydispersity and semiflexibility impact on the translational diffusion coefficient and electrophoretic mobility properties of actin filaments. An asymmetric, exponential length distribution for hydrodynamic conditions is used to characterize the actin filament polydispersity and the disparate rate lengths of barbed and pointed ends. Additionally, a modified cylindrical wormlike chain model was used to characterize the filament semiflexibility, effective monomer charge and diameter. The resulting expressions for the polydisperse translational diffusion coefficient and electrophoretic mobility depend on the persistence length, the effective filament diameter, the monomer charge, and the increasing rate length of barbed and pointed ends of the filaments. We considered typical experimental values for the degree of polymerization (370 G-actin proteins per um) and associate rates (barbed end ten times larger than the pointed end). The values for the other parameters were adjusted to reproduce the experimental data obtained for three typical polymerization buffers. This characterization is innovative since these parameter values are obtained from non-invasive experiments, and using the same experimental and hydrodynamic conditions.

Although buffers #1 and #2 produced some similar polyelectrolyte and hydrodynamic properties of actin filaments, many parameters account for the theoretical differences, such as the elongation rates. Nevertheless, buffer #3 displayed substantial differences in the actin structural conformations. Compared to those values obtained from molecular structure models, our results revealed a lower value of the effective G-actin charge and a more significant value of the effective filament diameter due to the formation of the double layer of the electrolyte surrounding the filaments. Additionally, compared to the values usually reported from electron micrographs, the lower values of our results for the persistence length and average contour filament length agrees with the significant difference in the association rates at the filament ends that shift to sub-micro lengths, the maximum of the length distribution. The polydispersity index ratio remains constant among the three buffers, which agrees with previous results. Buffer #3 revealed the formation of much longer filaments and bundles compared to the other two buffers caused by an increase in the association rates coming from the 2.5 times higher DTT concentration in the chemical composition. Buffer #3 also showed a lower translational diffusion coefficient, suggesting that actin filaments in this buffer were formed longer in average. Unlike buffer #2, the shearing parameter for buffer #3 resembles buffer #1’s result, where the filaments are more commonly fractioned due to the longer average filament lengths. This revealed an inverse proportionality correlation between the shearing parameter and the DTT concentration in the solution. We also analyzed the polarizability parameter, where the value for buffer #3 resulted higher than those in buffers #1 and #2. As the filament becomes more elongated in buffer #3, more charges/electrons are easily moved within the e-cloud/layers, increasing their polarizability and strengthening the dispersion forces, unlike compact molecules where all charges are symmetrically together. As a result, the formation of longer filaments generates higher polarizability values. Likewise, the value of the radius of gyration for buffer #3 was larger than those in buffers #1 and #2. From the comparison of the values of these parameters for rigid and semiflexibilty models, we concluded that actin filament semiflexibility contributes ~ 20% in the value of these parameters.

The optimized models and theories obtained in this article can be used and extended to calculate other actin filament’s properties, including stability, the intrinsic viscosity[9], molecular weight (Mark-Houwink exponential coefficient), the axial tension, the elastic stretch modulus [55], and the force-extension associated with the growth in length or the compression on the filament’s shrinkage[56]. Additionally, the fitting and optimization approaches described in this article can be used with other buffers, electrolyte conditions, and polydisperse charged semiflexible biopolymers.

## Author Contributions

Conceptualization, EA, MM and LB; methodology, EA and MM; validation, EA and MM; formal analysis, EA, AG, LB, and MM; investigation, EA and MM; experimental work, EA and AG; resources, MM; writing-original draft preparation, EA; writing-review and editing, MM and LB; supervision, MM; project administration, MM; funding acquisition, MM. All authors have read and agreed to the published version of the manuscript.

## Acknowledgements

We thanks Dr. Arturo Ponce for his support in the microscopy sample preparation and data analysis.

## Institutional Review Board Statement

This study did not require ethical approval; all data are available in the public domain.

## Informed Consent Statement

Not applicable.

## Data Availability Statement

Some or all data, models, experiments, or code that support the findings of this study are available from the corresponding author upon resonable request.

